# Microbiome Composition Regulates Cathepsin Expression in Vaginal Fluid Across BV Diagnoses and Community State Types

**DOI:** 10.64898/2026.05.07.723359

**Authors:** Carleigh Coffin Sokolik, Kayla Sahadeo, Jacqueline Vyce, Melissa Thomas, Cameron Celeste, Wambui Gachunga, Thaisha Calixte, Irene Ledford, Ji’Reh Williams, Erica Estess, Catera Wilder, Ivana K. Parker

## Abstract

**Purpose:** Bacterial vaginosis (BV) is associated with disruption of the vaginal microbiome and extracellular matrix (ECM) remodeling, yet the contribution of host proteases to this process remains unclear. This study investigated whether expression and activity of cathepsins K, L, S, and V differ by BV diagnosis and community state type (CST). We hypothesized that BV and BV associated CSTs would exhibit increased expression and activity of collagen and elastin-degrading cathepsins.

**Methods:** Vaginal fluid samples were collected and classified by BV diagnosis and CST. Cathepsin expression was evaluated by Western blotting to distinguish inactive and active enzyme forms. Proteolytic activity was assessed using multiplex cathepsin zymography. Statistical analyses compared cathepsin expression and activity across diagnoses and CSTs. Principal component analysis and linear regression were performed to assess associations between cathepsin activity, microbial diversity, and CST.

**Results:** Procathepsin K expression was significantly increased in BV-positive and CST IV samples, while total cathepsin L expression was significantly elevated in samples with Nugent-intermediate scores. Cathepsins S and V showed variation in inactive and active forms in Nugent-intermediate and CST III samples. In contrast, total cathepsin activity, including cathepsins K and V, did not significantly differ across BV diagnoses or CSTs. Overall, cathepsin activity varied between individuals rather than by clinical classification.

**Conclusions:** Cathepsin expression and maturation state differ by microbiome composition, suggesting that the vaginal microbiome may regulate post-translational processing of cathepsins. As a result, cathepsin activity appears to be regulated at the individual level rather than strictly by BV diagnosis or CST. These findings link vaginal microbiome composition to ECM remodeling and potential adverse reproductive outcomes.

## INTRODUCTION

The vaginal microbiome is a dynamic community of microorganisms that play a central role in gynecological and reproductive health. Bacterial vaginosis (BV) is a highly prevalent vaginal condition characterized by a shift from a *Lactobacillus-*dominated community to an anaerobic, polymicrobial microbiome [1], [2]. A healthy vaginal microbiome is typically dominated by *Lactobacillus* species, which maintain a low vaginal pH through lactic acid production, generate antimicrobial compounds, and reinforce epithelial barrier integrity. Depletion of these protective species has been associated with an increased risk of infection and adverse reproductive health outcomes [3], [4], [5], [6].

BV can be diagnosed syndromically using Amsel’s criteria, requiring at least three of the following: vaginal pH>4.5, presence of clue cells, thin grey/white milky discharge, and a positive whiff test [7]. BV can also be diagnosed using Nugent scoring, a Gram stain-based method performed on a vaginal smear that quantifies morphotypes associated with *Lactobacillus* depletion and anaerobic overgrowth [8]. Importantly, BV is associated with increased inflammation, altered immune responses, epithelial barrier disruption, and enhanced mucosal permeability [9], [10], [11], [12], [13], [14], [15]. Clinically, BV is linked to a heightened risk of STI and HIV acquisition and transmission, adverse pregnancy outcomes, infertility, and gynecological malignancies [16], [17], [18], [19], [20]. Given its high prevalence, recurrence rates, and systemic reproductive consequences, understanding the mechanistic pathways linking BV-associated dysbiosis to tissue dysfunction is critical.

Cathepsins are lysosomal cysteine proteolytic enzymes that regulate extracellular matrix (ECM) degradation. Several cathepsins, including K, L, S, and V, are capable of degrading structural ECM components such as collagen, elastin, laminin, and fibronectin [21], [22], [23]. Cathepsins are synthesized as inactive preproenzymes and undergo proteolytic removal of their prodomain within acidic endosomal compartments to generate mature enzymes [24], [25], [26], [27]. Activity of cathepsins is regulated by transcriptional control, pH, presence of inhibitors, and post-translational modifications [28], [29], [30], [31], [32], [33], [34], [35]. Under inflammatory conditions, cathepsins can be secreted extracellularly, where they can contribute to pathogenesis through aberrant ECM degradation, immune cell infiltration, and inflammation [36], [37], [38], [39]. Upregulated cathepsin activity has been implicated in a range of disease states including cancer, cardiovascular disease, and neurodegenerative disorders [40], [41], [42], [43], [44], [45], [46], [47]. In disease, aberrant cathepsin activity promotes cytokine release, activates additional proteases, alters cellular phenotypes, and disrupts ECM organization and stiffness [48].

Vaginal microbiome composition can also influence tissue remodeling processes by modulating protease expression and activity [49], [50]. In the vagina, ECM provides structural support and can control cell signaling and behavior [51]. Controlled ECM remodeling is essential for maintaining normal tissue homeostasis; however, microbes can secrete ECM-degrading enzymes which can alter matrix structure and mechanical properties, disrupting cellular signaling and promoting pathological host responses [52], [53], [54], [55]. In BV, increased inflammation and microbial-derived enzymes, including matrix metalloproteinases (MMPs), promote host protease activation, leading to collagen and elastin degradation, altered tissue stiffness, and barrier dysfunction [49], [50]. These structural changes may facilitate adverse health outcomes, thus, understanding how microbiome composition influences ECM remodeling providing mechanistic insight into how dysbiosis influences disease risk.

Advances in sequencing technologies have allowed for deeper characterization and identification of microbes within vaginal fluid. Given the complexity of the vaginal microbiome, vaginal microbial communities are often categorized into community state types (CSTs), characterized by the dominant microbial groups. CST I is dominated by *Lactobacillus crispatus*, CST II by *L. gasseri*, CST III by *L. iners*, CST V by *L. jensenii*, and CST IV is characterized by a diverse, non-*Lactobacillus*-dominated microbial community. Although CST III microbiomes are dominated by a *Lactobacillus* species, they are less stable and frequently transition to the more diverse CST IV state [56], [57]. CSTs may also influence protease expression which has not been explored.

Previous studies have shown increased expression and activity of MMPs in BV vaginal fluid; however, the impact of vaginal microbiome composition (CST) on cathepsin expression remains unexplored, despite cathepsins being more potent proteases [49]. In this study, we investigated whether BV diagnosis and CST-defined microbiome composition are associated with both altered expression and activity of cathepsins K, L, S, and V. Studying both cathepsin expression and activity is essential to determine whether increased enzyme abundance in BV-associated disease states translates into functional proteolytic activity that drives irregular ECM remodeling and inflammation. We hypothesize that vaginal fluid from women with BV will exhibit an increase in collagen- and elastin-degrading cathepsins, providing a potential mechanism for aberrant ECM remodeling and disruption of the epithelial and mucosal barriers observed in BV-associated disease.

## METHODS

### Ethical approval

This study was conducted under approval from the University of Florida Institutional Review Board (IRB202200874). Vaginal fluid samples were collected by a licensed nurse practitioner at the University of Florida Student Health Center in Gainesville, Florida. All participants provided informed consent prior to sample collection, and all procedures adhered to relevant institutional and federal guidelines for research involving human subjects.

### Sample collection

Vaginal fluid was collected by a nurse practitioner at the University of Florida Student Health Center in Gainesville, FL using a nylon flocked swab (FLOQ Swab 519C) (Table 1). A positive BV diagnosis was made using Amsel’s criteria if at least three of the following were present: positive whiff test, thin white/grey homogeneous discharge, vaginal pH > 4.5, and presence of clue cells. In parallel, vaginal smears were Gram-stained and evaluated using the Nugent scoring system to provide a standardized, microscopy-based assessment of BV. Slides were examined under light microscopy to characterize bacteria morphotypes, including gram-positive rods, gram-variable rods, and curved gram-negative rods. Scores were assigned on a 0-10 scale according to established criteria where 0-3 indicated Lactobacillus dominant microbiota (BV-negative), 4-6 indicated intermediate microbiota, and 7-10 indicated BV (BV-positive) [8]. Additional swabs were collected at the time of visit and the vaginal microbiome was characterized for each individual using metagenomic next generation sequencing performed by Evvy.

**Table 1.** Cohort Description.

|  |  | Participants (n = 19) |
| --- | --- | --- |
| Age | 18-30 | 15 |
|  | 31-40 | 2 |
|  | 41-50 | 1 |
|  | 51-60 | 1 |
| Race | Asian | 4 |
|  | Black or African American | 3 |
|  | Hispanic or Latino | 8 |
|  | White | 13 |
|  | Other | 1 |
| Nugent Score | 0-3 | 5 |
|  | 4-6 | 6 |
|  | 7-10 | 6 |
| Amsel Diagnosis | Negative | 13 |
|  | Equivocal | 0 |
|  | Positive | 6 |
| Community State Type | I | 6 |
|  | II | 0 |
|  | III | 6 |
|  | IV | 7 |
|  | V | 0 |
| Vaginal pH Category | Low (pH < 4.5) | 5 |
|  | High (pH ≥ 4.5) | 14 |

Participants were included in the study if they were >18 years of age and provided informed consent. Participants were excluded from the study if they were pregnant at time of study, had known STIs/HIV, had vaginal intercourse within 1 day of collection, or had their menstrual cycle within 3 days of collection.

### Sample processing and storage

After collection, flocked swabs and Evvy swabs were stored at -80°C. Evvy Vaginal Health Test kits were shipped to Evvy within one month of collection. Vaginal fluid was isolated from flocked swabs by adding 750 μL of zymogram lysis buffer with 0.1 mM leupeptin to the dry tube [58]. Samples were sonicated with two pulses at 20% amplitude on ice before centrifuging at 10,000 x g for 5 minutes. The supernatant was divided into 125 μL aliquots and stored at -80C.

### Micro BCA Assay

To measure the total protein concentration within each sample, the Pierce Micro BCA Protein Assay Kit was used. Protein concentration was determined by comparing absorbance values of our samples to a standard curve of bovine serum albumin at known protein concentrations. The R^2^ value of the standard curve needed to be ≥ 0.99 to be accepted.

### Multiplex Cathepsin Zymography

To detect amounts of active cathepsins K, L, S, and V, multiplex cathepsin zymography was performed using a previously established protocol [58]. Substrate gel electrophoresis was performed by loading 15 μg of protein from each sample into each well at a total volume of 18.8 μL. Additionally, 18.8 μL of prestained molecular weight standard (SKU: 08W00027) was added at each end of the gel as a protein ladder, and 18.8 μL of 1X sample nonreducing loading buffer was added as a negative control. Electrophoresis was performed at 200 V for 1 hour.

After electrophoresis, polyacrylamide gels were removed and placed into containers with renaturing buffer for three 10-minute rinses to renature enzymes and cleave substrate proteins. Gels were then equilibrated in 25 mL pH 6 assay buffer for 30 minutes, before incubating in fresh 25 mL of assay buffer overnight at 37 °C. After a 24 hr incubation period, gels were stained in Coomassie blue for 1 hr, and destained in destain buffer for 1 hr. The presence of white bands on a dark blue background indicates the presence of cathepsin activity.

Gels were imaged on Amersham ImageQuant 800 GxP biomolecular imager using the fluorescence setting at 680 nm to detect Coomassie blue and 780 nm to detect the prestained molecular weight standard. Densitometry of the signals was performed using the ImageQuant TL analysis software.

### Western Blotting

Western blotting for Cathepsins K, L, S, and V was repeated in three independent experiments for each sample. 15 μg of protein per sample was combined with 1 μL of Amersham WB Cy5 dye, and 5X reducing loading buffer (50 mM Tris-HCl pH 6.8, 8% glycerol, 16% SDS, 4% beta-mercaptoethanol, 0.04% bromophenol blue) to reach a total volume of 18.8 μL to add to each well. Each sample was boiled for 5 minutes at 95 °C and centrifuged at 16,000 x g for 1 minute. Samples were then loaded onto a 12% SDS-page gel along with a prestained protein ladder on each end of the gel, and 1X reducing loading buffer as a negative control.

Electrophoresis was performed at 155 V for 75 minutes. Proteins were transferred from the gel to a PVDF card (Immobilon-P, Millipore) using a semi-dry Western blot transfer protocol. Membranes were blocked with blocking buffer made of 3% BSA diluted in TBST (20 mM Tris-HCl pH 7.5, 150 mM NaCl, 0.1% Tween-20) for 1 hour at room temperature. Primary antibodies were prepared in blocking buffer to incubate membranes overnight at 4 °C. Primary antibodies against Cathepsin K (#11239-1-AP) and Cathepsin L (#27952-1-AP) were purchased from Proteintech, and Cathepsin S (#PA5-30002) and Cathepsin V (#PA5-112393) were purchased from Invitrogen. Cy5-prelabeled total protein served as a loading control.

After overnight incubation with primary antibodies, membranes were washed five times with TBST, then incubated with a secondary antibody (Amersham ECL Rabbit IgG, HRP-linked whole Ab, Cytiva # NA934) diluted 1:10,000 with 1X PBS for 1 hour at room temperature. Membranes were washed once again five times with TBST, then incubated for 1 minute with Amersham ECL detection reagents (Cytiva) and imaged in an Amersham ImageQuant 800 GxP biomolecular imager using chemiluminescence settings to detect HRP-tagged secondary antibodies and fluorescence settings to detect Cy5 labeled total protein. Densitometry of the signals was performed using ImageQuant TL analysis software. The background was subtracted from the target protein bands and Cy5-labeled total protein with default rolling ball sizes.

### Statistical Analysis

To compare measures between two groups either an unpaired t-test or Mann-Whitney U test was used, depending on whether the measures were normally distributed. Shapiro-Wilk test was used to determine normality of data sets. To compare measures between three groups either a one-way ANOVA or Kruskal-Wallis test was used, depending on the normality of the data sets.

## RESULTS

### Identifying CSTs and microbial diversity within samples

To evaluate the relationship between vaginal microbiome composition and cathepsin expression and activity, a cohort of 19 participants with and without BV was established (Table 1). BV status was assessed using both Amsel criteria and Nugent scoring. Using Amsel criteria, 13 participants received a negative BV diagnosis, 0 received an equivocal diagnosis, and 6 received a positive diagnosis (Table 1). For Nugent scoring, 5 samples had BV-negative scores between 0-3, 6 samples had BV-intermediate scores between 4-6, and 6 samples had BV-positive scores between 7-10 (Table 1). Vaginal microbiome composition was characterized using metagenomic next-generation sequencing to quantify the relative abundances of 445 microbial species.

Principal component analysis (PCA) of species-level relative abundance data revealed clustering of samples by Amsel diagnosis, Nugent score, and CST. The first principal component (PC1) was primarily associated with *Lactobacillus crispatus*, while the second principal component (PC2) was associated with *Lactobacillus iners* (Figure 1A-C). From PCA, 6 samples were identified as CST I, 6 samples were identified as CST III, and 7 samples were identified as CST IV (Table 1 and Figure 1C). To quantify within-sample microbial diversity, the Shannon Diversity Index (SDI) was calculated for each sample, with higher SDI values indicating greater microbial diversity.

**Fig. 1.**
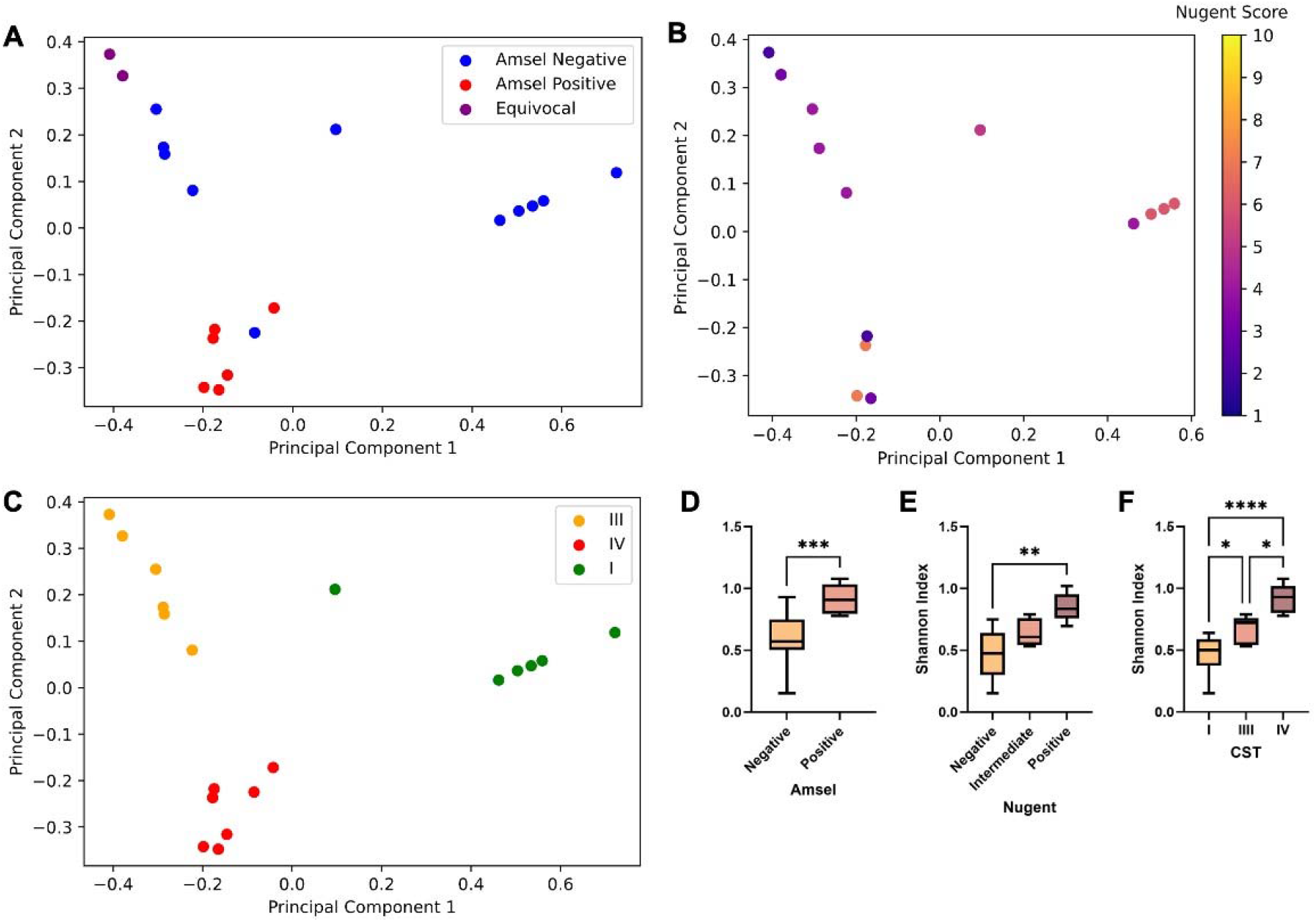
Vaginal microbiome composition and microbial diversity of samples. PCA of species-level relative abundance data with legends by **A** Amsel diagnosis, **B** Nugent diagnosis, and **C** CST. PC1 is primarily associated with *Lactobacillus crispatus* and PC2 with *Lactobacillus iners*. SDI was calculated for each of the samples and is shown by **D** Amsel diagnosis, **E** Nugent diagnosis, and **F** CST. Statistical significance denoted by *p≤0.05, **p≤0.01, ***p≤0.001, ****p≤0.0001.

SDI values were significantly higher in samples from participants with a positive Amsel diagnosis compared to those with a negative (p < 0.001) diagnosis (Figure 1D). Similarly, samples classified with a positive Nugent score exhibited significantly higher SDI values compared to Nugent-negative samples (p < 0.01) (Figure 1E). SDI values also varied significantly by CST, with CST IV microbiomes exhibiting significantly higher SDI values compared to CST I (p < 0.0001) or CST III (p < 0.05) samples (Figure 1F). Additionally, CST III samples had significantly higher SDI values than CST I samples (p < 0.05) (Figure 1F).

### Cathepsin Activity Across Vaginal Microbiome Classifications

To investigate how the vaginal microbiome composition affects cathepsin activity, cathepsin K (37 kDa) and cathepsin V (35 kDa) were examined using cathepsin zymography (Figure 2B-J). Overall significant differences (p < 0.05) in cathepsin activity were not observed across Amsel diagnostic categories, Nugent score classifications, or CSTs (Figure 2 B-J).

**Fig. 2.**
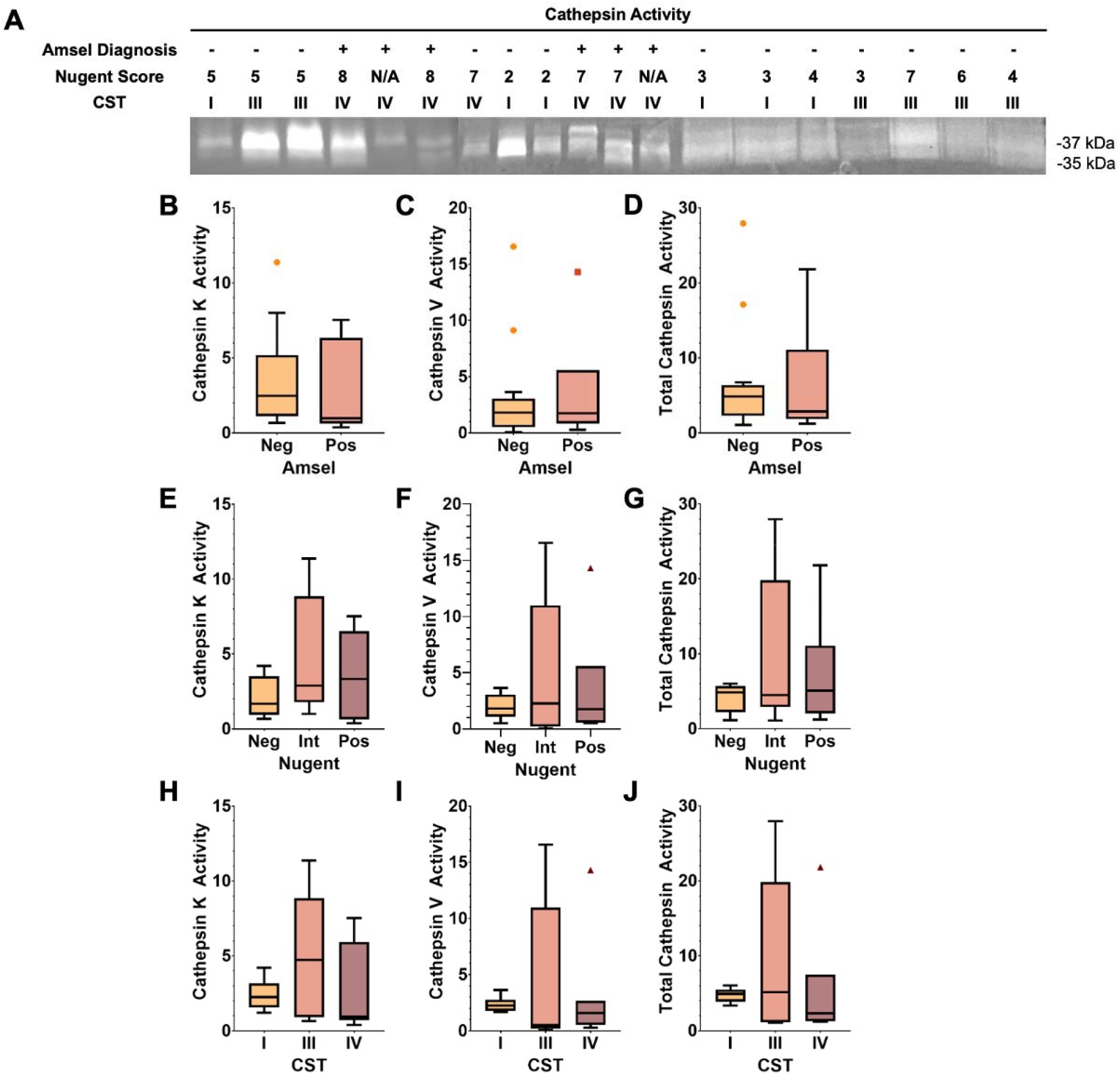
Cathepsin activity in vaginal fluid across vaginal microbiome classification. **A** Representative cathepsin zymography from 19 patients stratified by Amsel diagnosis, Nugent score, and CST; cathepsin K (∼37 kDa) and V (∼35 kDa) are shown. By Amsel diagnosis, quantified **B** cathepsin K, **C** cathepsin V, and **D** total activity are shown; by Nugent score **E** cathepsin K, **F** cathepsin V, and **G** total activity; and by CST, **H** cathepsin K, **I** cathepsin V, and **J** total activity.

### Cathepsin K and L expression is increased in non-optimal vaginal microbiomes

For cathepsins to degrade ECM, inactive pro-cathepsins must be enzymatically cleaved to reveal the active site. Western blotting was performed and differences in the inactive precursor and mature active forms of potent collagenases cathepsins K and L were determined.

Procathepsin K (∼47 kDa) and mature cathepsin K (∼27 kDa) were detected in vaginal fluid samples (Figure 3A). Procathepsin L (∼45 kDa) and mature cathepsin L (∼25 kDa) were similarly identified (Figure 4A). Total cathepsin expression was calculated as the sum of quantified precursor and mature band intensities.

**Fig. 3.**
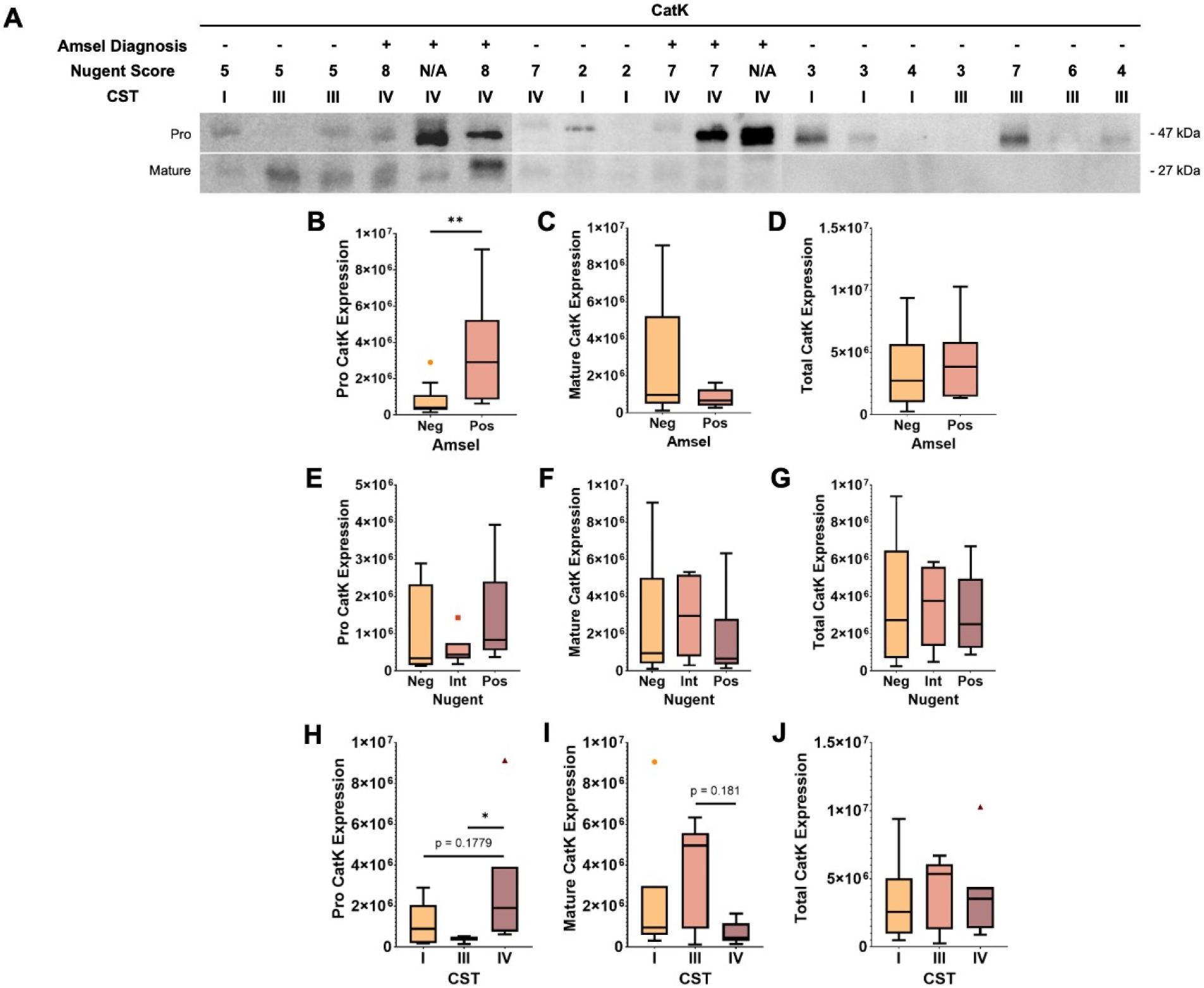
Cathepsin K expression in vaginal fluid across vaginal microbiome classification. **A** Representative cathepsin K western blot from 19 patients stratified by Amsel diagnosis, Nugent score, and CST; pro cathepsin K (∼47 kDa) and mature cathepsin K (∼27 kDa) are shown. By Amsel diagnosis, quantified **B** pro cathepsin K, **C** mature cathepsin K, and **D** total expression are shown; by Nugent score **E** pro cathepsin K, **F** mature cathepsin K, and **G** total expression; and by CST, **H** pro cathepsin K, **I** mature cathepsin K, and **J** total expression. Statistical significance denoted by *p≤0.05, **p≤0.01.

**Fig. 4.**
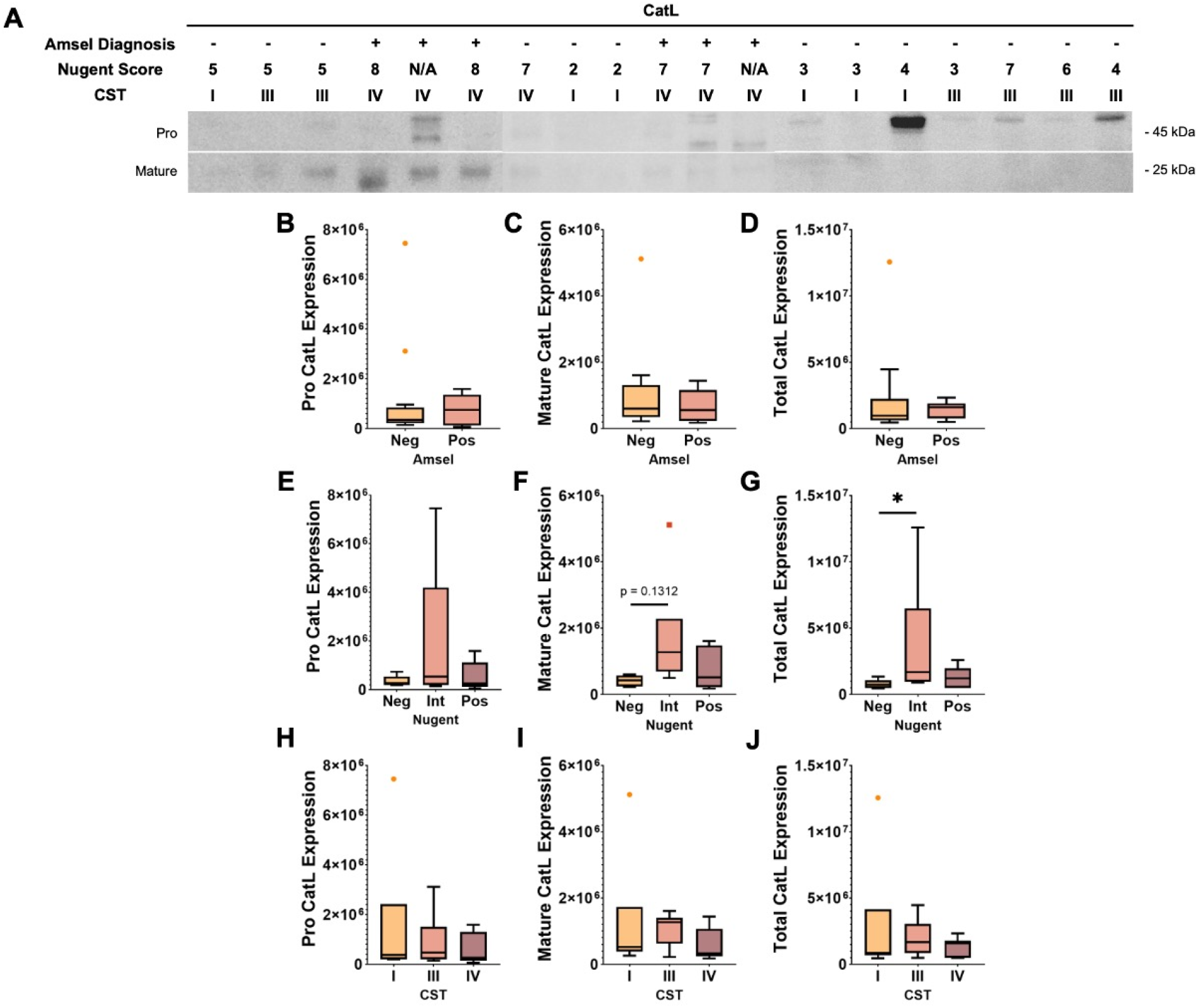
Cathepsin L expression in vaginal fluid across vaginal microbiome classification. **A** Representative cathepsin L western blot from 19 patients stratified by Amsel diagnosis, Nugent score, and CST; pro cathepsin L (∼45 kDa) and mature cathepsin L (∼25 kDa) are shown. By Amsel diagnosis, quantified **B** pro cathepsin L, **C** mature cathepsin L, and **D** total expression are shown; by Nugent score **E** pro cathepsin L, **F** mature cathepsin L, and **G** total expression; and by CST, **H** pro cathepsin L, **I** mature cathepsin L, and **J** total expression. Statistical significance denoted by *p≤0.05.

Procathepsin K expression was significantly increased in samples from participants with a positive Amsel diagnosis compared to BV-negative participants (P < 0.01) (Figure 3B) and in CST IV relative to CST III (p < 0.05) (Figure 3H). Although not statistically significant, procathepsin K levels were also higher in CST IV compared to CST I (p = 0.1779) (Figure 3H). Mature cathepsin K expression was elevated in CST III relative to CST IV (p = 0.1810) (Figure 3I).

For cathepsin L, total expression was significantly increased in samples with intermediate Nugent scores compared to Nugent-negative samples (p < 0.05) (Figure 4G). Mature cathepsin L expression also trended higher in intermediate Nugent samples relative to Nugent-negative samples (p = 0.1312) (Figure 4F).

Together, these findings suggest that BV-associated microbiome states, particularly Amsel-positive, CST III, and CST IV classifications, are associated with increased cathepsin K expression, indicating potential dysregulation of proteolytic activation. Increased total and mature L expression in Nugent-intermediate samples further supports a shift in protease abundances across vaginal microbiome states that may influence ECM remodeling.

### Cathepsin S and V Expression in Vaginal Fluid

Western blotting was performed to quantify precursor and mature forms of cathepsins S and V, elastases implicated in ECM remodeling in vaginal fluid with different BV diagnoses and microbiome compositions.

For cathepsin S, pre-pro-cathepsin (∼50 kDa), procathepsin (∼40 kDa), a single-chain mature form (∼35 kDa), and mature heavy (∼25 kDa) and light (∼10 kDa) chains were detected (Figure 5A). Preprocathepsin S expression was significantly increased in samples with intermediate Nugent scores compared to Nugent-negative samples (p < 0.05) (Figure 5G) and showed similar upward trends relative to Nugent-positive samples (p = 0.0968) (Figure 5G). Additionally, samples from Amsel-positive and CST III samples had modestly higher preprocathepsin S expression than Amsel-negative (p = 0.21) (Figure 5B) or CST I (p = 0.2434) (Figure 5L) samples, respectively. Procathepsin S expression was also elevated in intermediate Nugent samples compared to Nugent-negative samples (p = 0.0662) (Figure 5H).

**Fig. 5.**
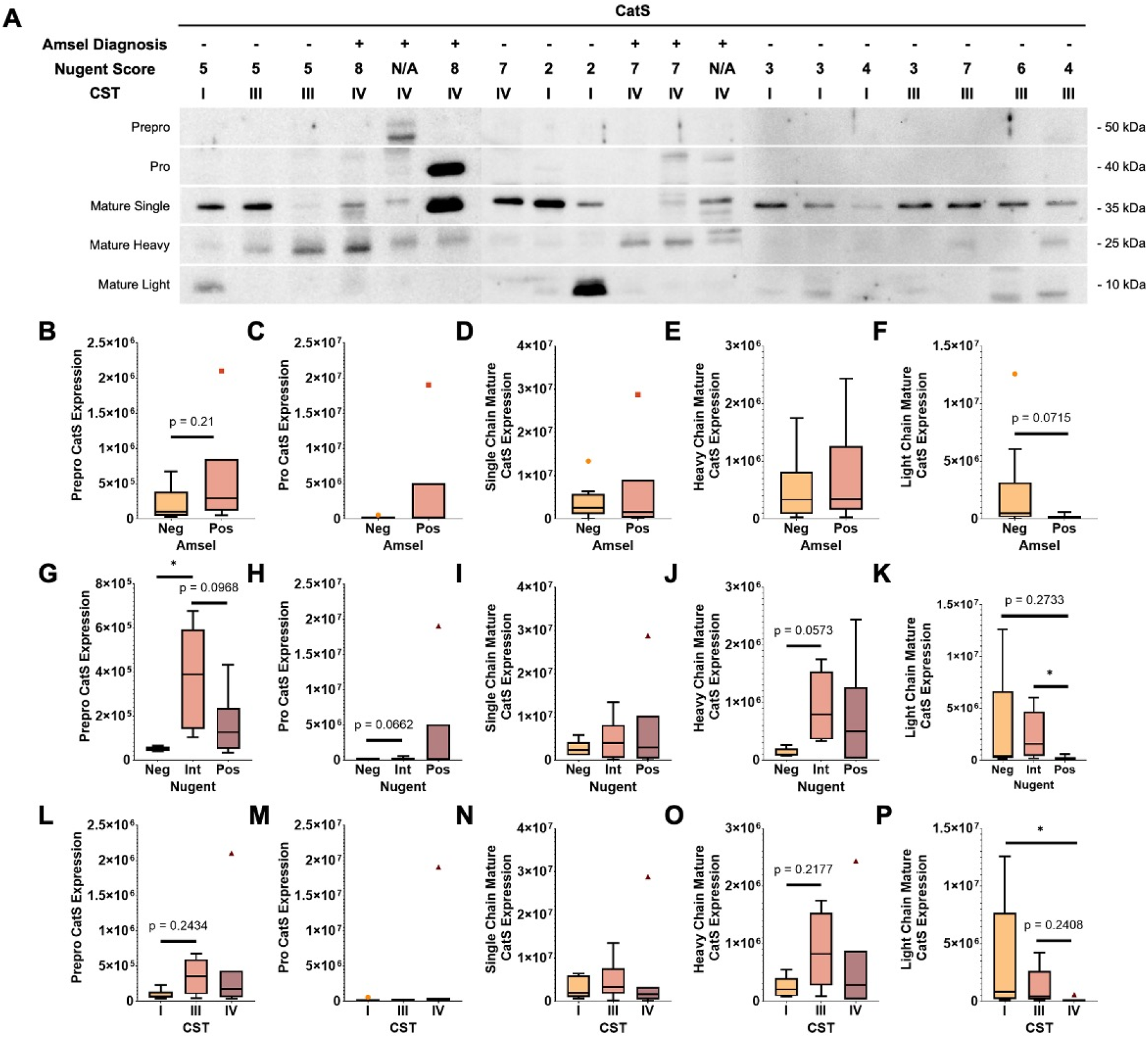
Cathepsin S expression in vaginal fluid across vaginal microbiome classification. **A** Representative cathepsin S western blot from 19 patients stratified by Amsel diagnosis, Nugent score, and CST; prepro (∼50 kDa), pro (∼40 kDa), mature single (∼35 kDa), mature heavy (∼25 kDa), and mature light (∼10 kDa) cathepsin S are shown. By Amsel diagnosis, quantified **B** prepro, **C** pro, **D** mature single, **E** mature heavy, and **F** mature light cathepsin S expression are shown; by Nugent score **G** prepro, **H** pro, **I** mature single, **J** mature heavy, and **K** mature single expression; and by CST, **L** prepro, **M** pro, **N** mature single, **O** mature heavy, and **P** mature light expression. Statistical significance denoted by *p≤0.05.

Mature cathepsin S light chain expression varied significantly by Nugent score and CST, with significantly higher levels in intermediate Nugent and CST I samples compared to Nugent-positive (p < 0.05) (Figure 5K) and CST IV samples (p < 0.05) (Figure 5P), respectively. Similarly, mature heavy chain expression showed increases in intermediate and CST III samples compared to Nugent-negative (p = 0.0573) (Figure 5J) and CST I samples (p = 0.2177) (Figure 5O), though not significant. Cathepsin S light chain expression was also higher in Nugent-negative and CST III samples in relation to Nugent-positive (p = 0.2733) (Figure 5K) and CST IV samples (p = 0.2408) (Figure 5P).

For cathepsin V, procathepsin (∼50 kDa), a single-chain mature form (∼37 kDa), and mature heavy (∼25 kDa) and light (∼10 kDa) chains were detected (Figure 6A). Procathepsin V expression was significantly higher in Amsel-positive samples compared to Amsel-negative samples (p < 0.05) (Figure 6B). Trends in procathepsin V elevations were also observed in Nugent-intermediate samples compared to samples with negative (p = 0.2372) and positive (p = 0.2822) scores (Figure 6F). Additionally, procathepsin V expression was higher in CST IV samples relative to CST I samples (p = 0.0662) (Figure 6J).

**Fig. 6.**
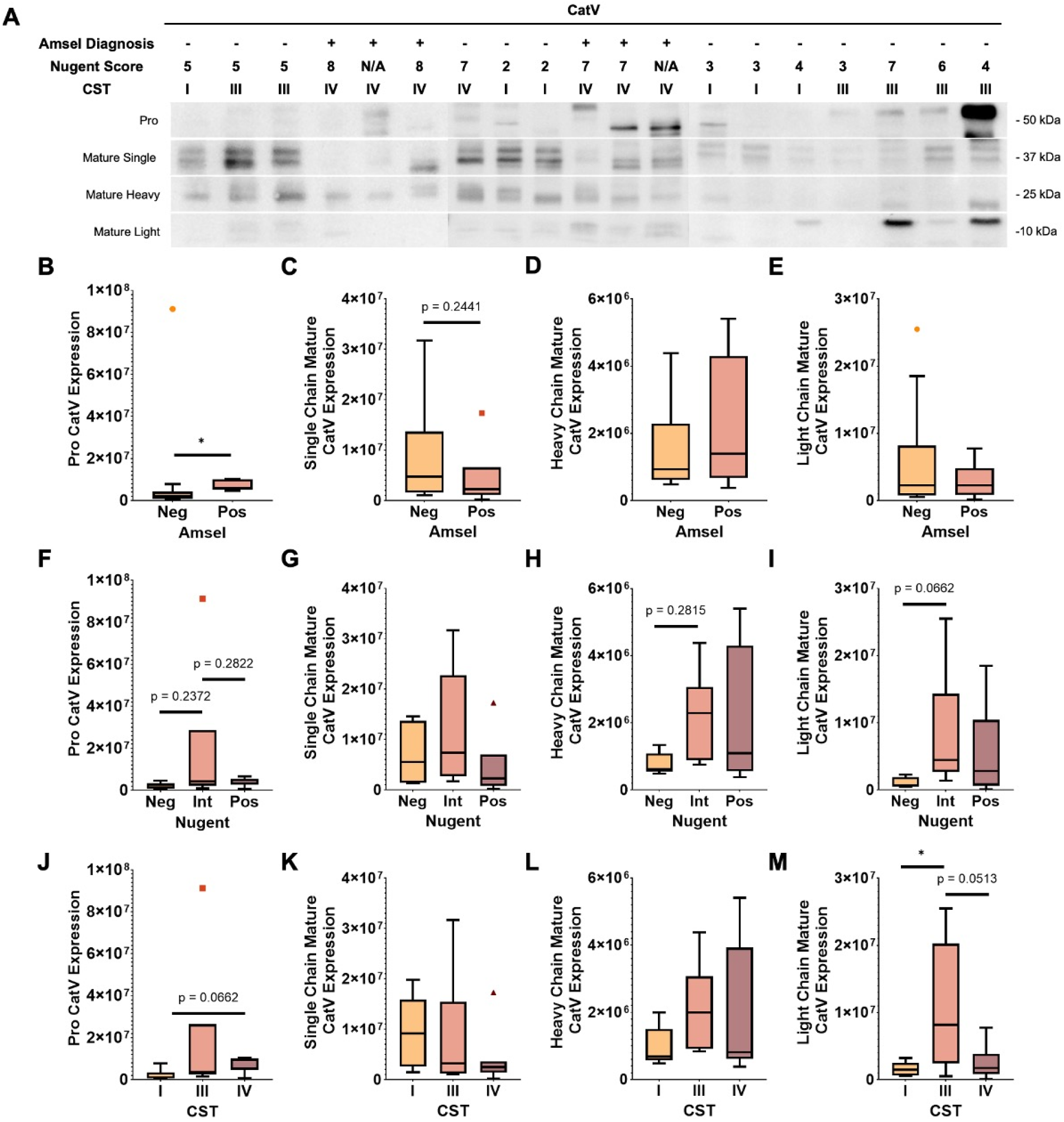
Cathepsin V expression in vaginal fluid across vaginal microbiome classification. **A** Representative cathepsin V western blot from 19 patients stratified by Amsel diagnosis, Nugent score, and CST; pro (∼50 kDa), mature single (∼37 kDa), mature heavy (∼25 kDa), and mature light (∼10 kDa) cathepsin V are shown. By Amsel diagnosis, quantified **B** prepro, **C** pro, **D** mature single, **E** mature heavy, and **F** mature light cathepsin V expression are shown; by Nugent score **G** prepro, **H** pro, **I** mature single, **J** mature heavy, and **K** mature single expression; and by CST, **L** prepro, **M** pro, **N** mature single, **O** mature heavy, and **P** mature light expression. Statistical significance denoted by *p≤ 0.05.

Single-chain mature cathepsin V expression was elevated in Amsel-negative samples compared to Amsel-positive samples, though this difference was not statistically significant (p = 0.2441) (Figure 6C). Expression of the mature heavy chain of cathepsin V was higher in Nugent-intermediate samples compared to Nugent-negative samples (p = 0.2815) (Figure 6H). Similarly, expression of the mature light chain was significantly increased in CST III samples relative to CST I samples (p < 0.05) (Figure 6M). Light chain expression was also higher in Nugent-intermediate and CST III samples compared to Nugent-negative (p = 0.0662) (Figure 6I) or CST IV (p = 0.0513) (Figure 6M) samples.

Collectively, these findings indicate that BV-associated and intermediate microbiome states are characterized by altered cathepsin S and V processing, with shifts in precursor and mature forms. Differences in expression of inactive and active elastases suggests that microbiome composition may influence cathepsin activation dynamics and ECM remodeling in the vaginal microenvironment.

## DISCUSSION

Defining the influence of the vaginal microbiome on tissue remodeling is critical to understand the mechanistic links between dysbiosis and adverse gynecologic and reproductive health outcomes. In this study, we sought to explore how BV or CST-defined vaginal microbiome composition influences the expression and activity of cathepsins, proteases implicated in ECM remodeling. Using western blotting and zymography, we detected cathepsin expression and activity in vaginal fluid samples, with a particular focus on cathepsins K, L, S, and V as they are potent collagenases and elastases. We observed that while overall cathepsin enzymatic activity did not differ by BV diagnosis or CST, cathepsin K, L, S, and V expressions were variable based on microbiome composition having distinct profiles associated with each maturation state.

Previous studies have reported significant increases in expression and activity of MMP-1, MMP-3, MMP-10, and MMP-13, proteases known to cleave ECM and tight junction proteins, in BV-positive cervicovaginal fluid. However, in our study multiplex cathepsin zymography revealed no statistically significant differences in cathepsin K, cathepsin V, or total cathepsin activity across Amsel or Nugent diagnoses, or CSTs. Previous studies used BioPlex immunoassays to quantify MMP concentrations and the SensoLyte 520 Generic MMP assay to measure overall enzymatic activity [49]. In contrast, our use of Western blotting and zymography enables detection of protein molecular weight, allowing us to distinguish inactive and active cathepsin forms within vaginal fluid samples. Although cathepsin activity did not vary by BV diagnosis or CST, individual patients did exhibit increased cathepsin activity compared to others suggesting that protease activation may be governed by patient-specific host-microbiome interactions rather than microbiome characterization alone. Additionally, because CSTs III and IV can be observed in both asymptomatic individuals and those with clinical BV, differences in cathepsin expression by CST may not translate to differences in activity, which is further regulated by post-translational events.

Western blot analyses revealed distinct differences in expression of cathepsin protein expression associated with microbiome composition. Notably, procathepsin K expression was significantly higher in Amsel-positive and CST IV samples, typically characterized by reduced *Lactobacillus* abundance and increased microbial diversity. Increased accumulation of the inactive precursor may indicate altered processing, secretion, or inhibition of proteases as a result of vaginal microbiome dysbiosis. Interestingly, expression of mature cathepsin K trended higher in CST III samples dominated by *Lactobacillus iners*. These findings indicate future studies with larger sample sizes should explore how vaginal microbiome composition affects collagen degradation, specifically through collagenase production.

Investigation of cathepsin L expression, another potent collagenase, further supported the role of intermediate vaginal microbiomes in ECM remodeling. Total cathepsin L expression was significantly increased in samples with an intermediate Nugent diagnosis, with a corresponding trend toward mature cathepsin L expression.

Differences in expression of cathepsins S and V, elastin degrading proteases, were also detected between BV diagnoses and CSTs. Inactive precursors preprocathepsin S and procathepsin V were elevated in samples with a positive Amsel diagnosis or intermediate Nugent diagnosis. Similarly, mature light and heavy chain forms of cathepsins S and V were more prominent in samples with intermediate Nugent scores or CST III microbiomes. These results indicate that increased production of elastin degrading cathepsins in intermediate vaginal microbiomes may modulate ECM remodeling processes and lead to a compromised epithelial barrier.

The modest sample size of this study limited statistical power, particularly given the high variability between vaginal fluid samples. Furthermore, CSTs are complex with recent studies showing distinct strains of the same species eliciting different environmental responses [59]. CST III, dominated by *Lactobacillus iners*, is a transitional state that can shift between both health and BV, limiting clear interpretation of its biological significance. Additionally, because BV diagnosis and CST are determined after sample collection, we are unable to control group sizes, making it difficult to obtain balanced representation across diagnoses and CSTs. The sampling variability in this study limits mechanistic conclusions but underscores the need for synthetic *in vitro* models that allow controlled manipulation of the vaginal microbiome to directly study microbiome-ECM interactions [60].

In conclusion, the results of this study suggest that the vaginal microbiome composition, particularly transitional states between health and dysbiosis, are associated with altered expression of cathepsin K, L, S, and V. Cathepsins are potent collagenases and elastases, playing an important role in ECM remodeling processes. These findings suggest that vaginal microbiome modulated cathepsin activity is a mechanism for adverse reproductive health outcomes.

## Consent to participate

All participants provided voluntary, written informed consent prior to enrollment in the study. Study aims, procedures, potential risks, and participant rights were explained by trained study staff, and participants were given the opportunity to ask questions before signing the consent form. Consent procedures were conducted in accordance with the University of Florida IRB-approved protocol (IRB202200874) and adhered to all institutional and federal guidelines for human subjects research.

## Consent to publish

All participants provided written consent permitting the use of their de-identified information for research dissemination. No identifiable personal information is included in this manuscript.

## Data Availability Statement

The data supporting the findings of this study include human participant information and are therefore subject to Institutional Review Board (IRB) restrictions. De-identified data generated and analyzed during the study are available from the corresponding author upon reasonable request and with approval from the University of Florida Institutional Review Board (IRB202200874). Identifiable participant data cannot be shared.

## Authors Contributions

C.C.S.: experimental design, data generation, data analysis, data interpretation and visualization, original manuscript drafting and revision. K.S., J.V., and M.T.: data generation, and data analysis. C.C.: data analysis. W.G., T.C.: Nugent scoring. I.L., J.W.: clinical coordination. E.E.: sample collection and diagnosing. C.W.: manuscript revision and editing. I.K.P. inception of study idea, data interpretation, manuscript revision and editing, supervision, and funding acquisition. Correspondence to Ivana K. Parker.

