## Supplemental Files for "Microbiome Composition Regulates Cathepsin Expression in Vaginal Fluid Across BV Diagnoses and Community State Types"

**SUPPLEMENTARY INFORMATION**

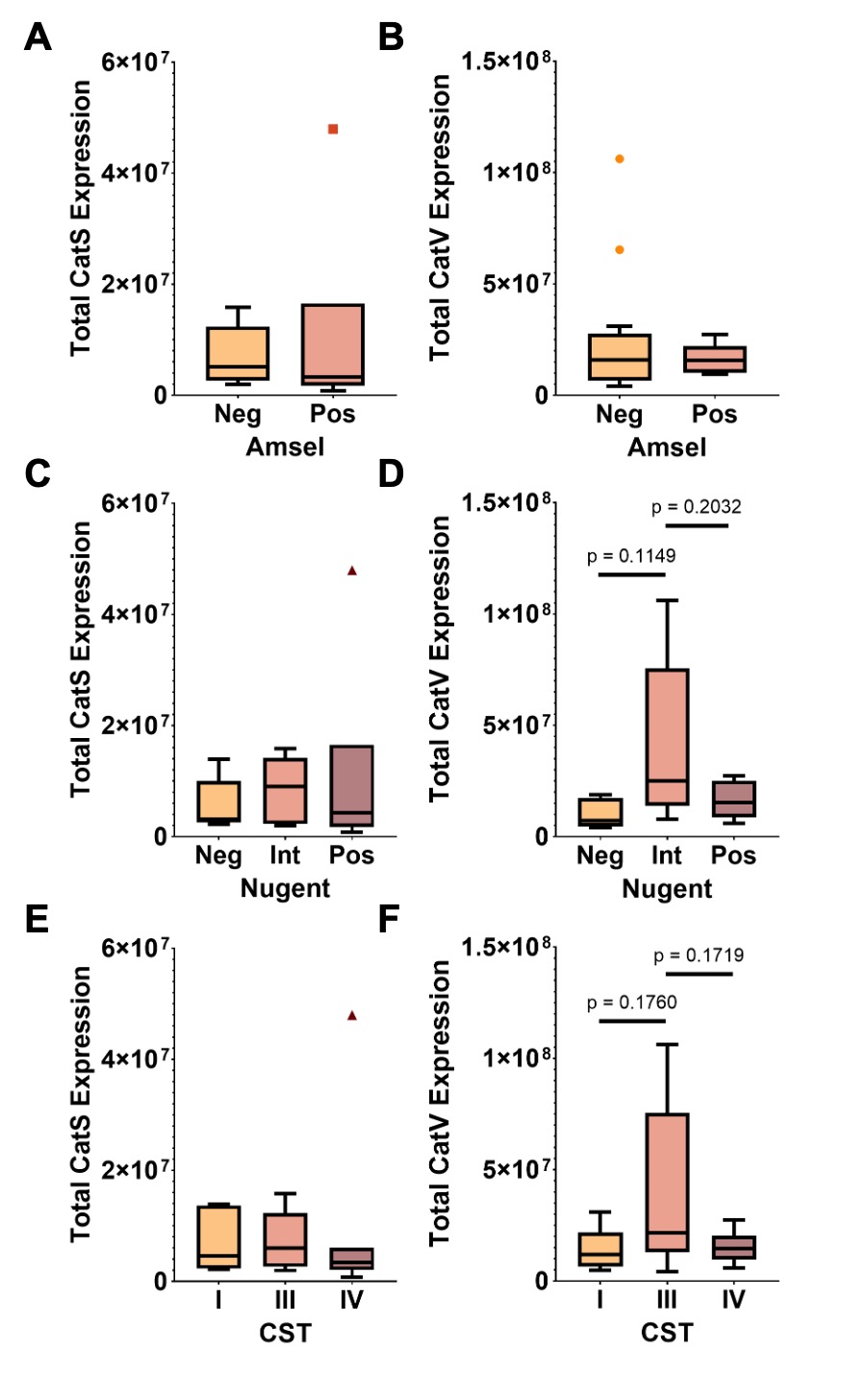

**Fig. S1** Total Cathepsin S and V expression in vaginal fluid across vaginal microbiome classification. By Amsel diagnosis, quantified total **A** cathepsin S, and **B** cathepsin V expression are shown; by Nugent score **C** cathepsin S, and **D** cathepsin V are shown; and by CST, **E** cathepsin S, and **F** cathepsin V are shown.

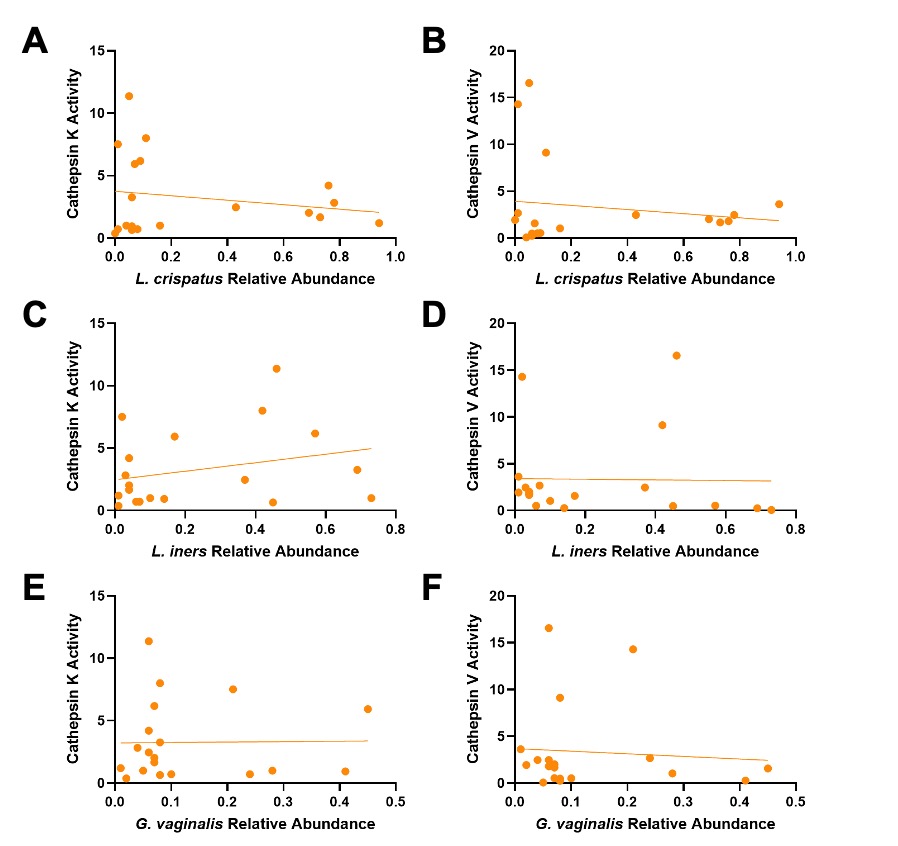

**Fig. S2** Cathepsin Activity regression analysis. By L. crispatus relative abundance, **A** cathepsin K, and **B** cathepsin V activity are shown; by L. iners relative abundance, **C** cathepsin K, and **D** cathepsin V are shown; and by G. vaginalis relative abundance, **E** cathepsin K, and **F** cathepsin V are shown.

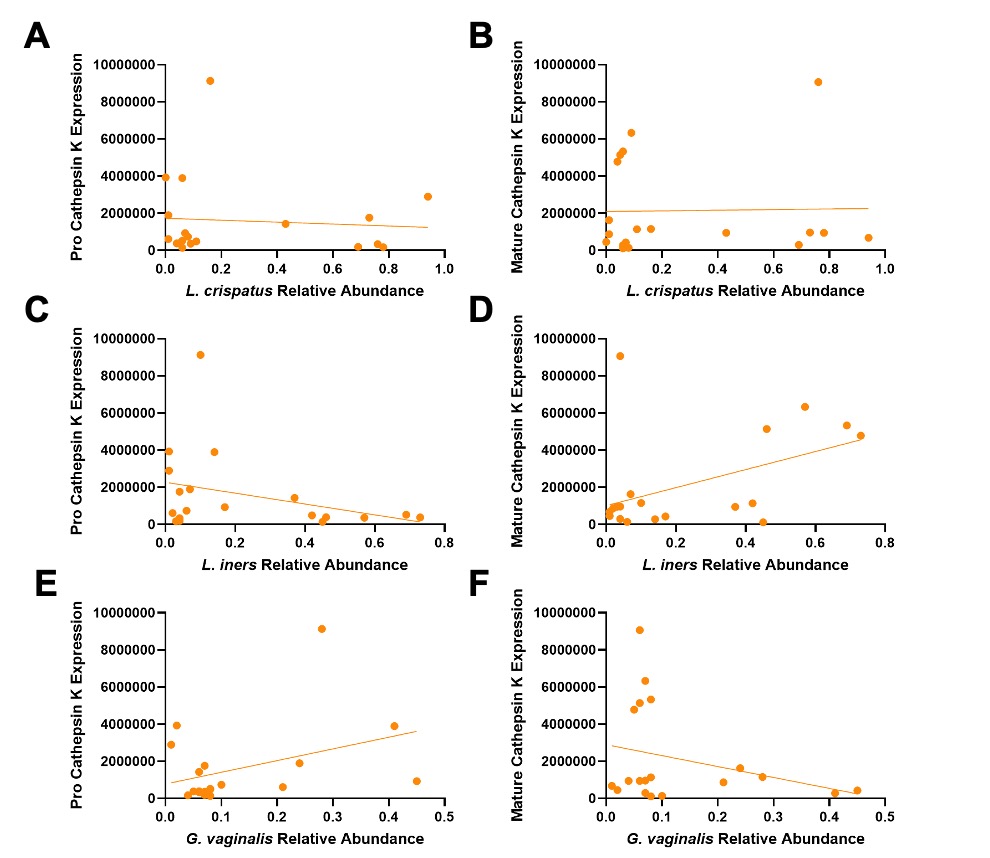

**Fig. S3** Cathepsin K expression regression analysis. By L. crispatus relative abundance, **A** pro, and **B** mature cathepsin K expression are shown; by L. iners relative abundance, **C** pro, and **D** mature cathepsin K are shown; and by G. vaginalis relative abundance, **E** pro, and **F** mature cathepsin K are shown.

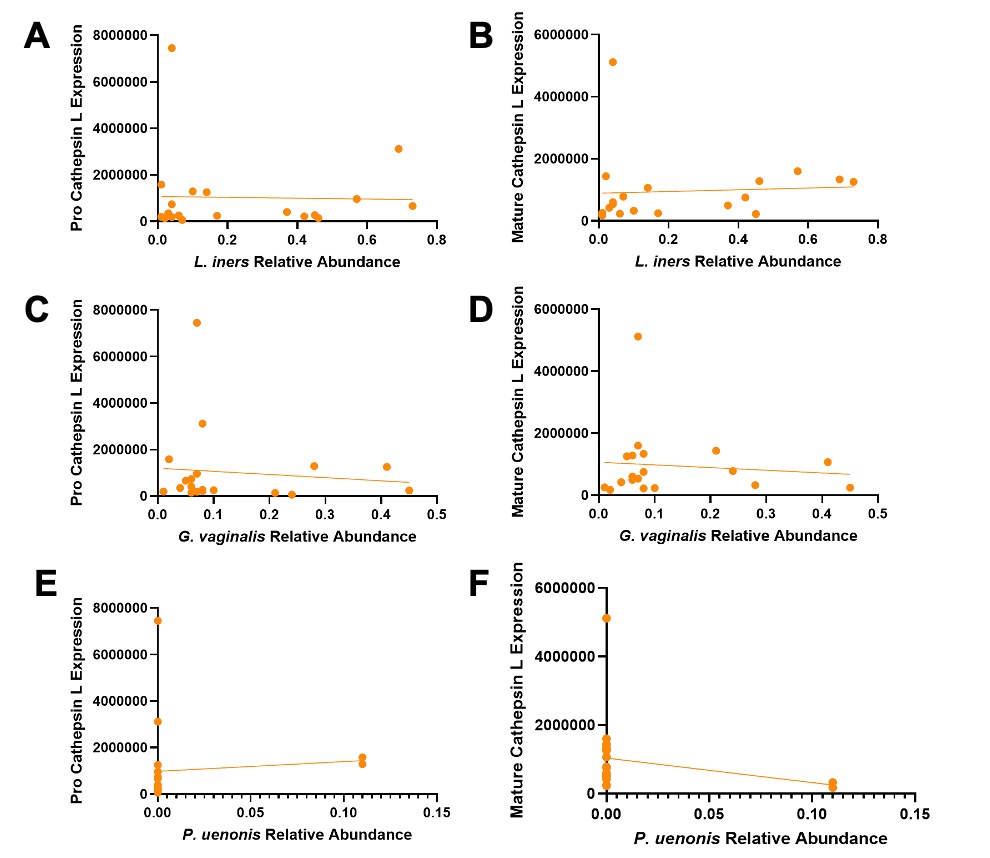

**Fig. S4** Cathepsin L expression regression analysis. By L. crispatus relative abundance, **A** pro, and **B** mature cathepsin L expression are shown; by L. iners relative abundance, **C** pro, and **D** mature cathepsin L are shown; and by G. vaginalis relative abundance, **E** pro, and **F** mature cathepsin L are shown.

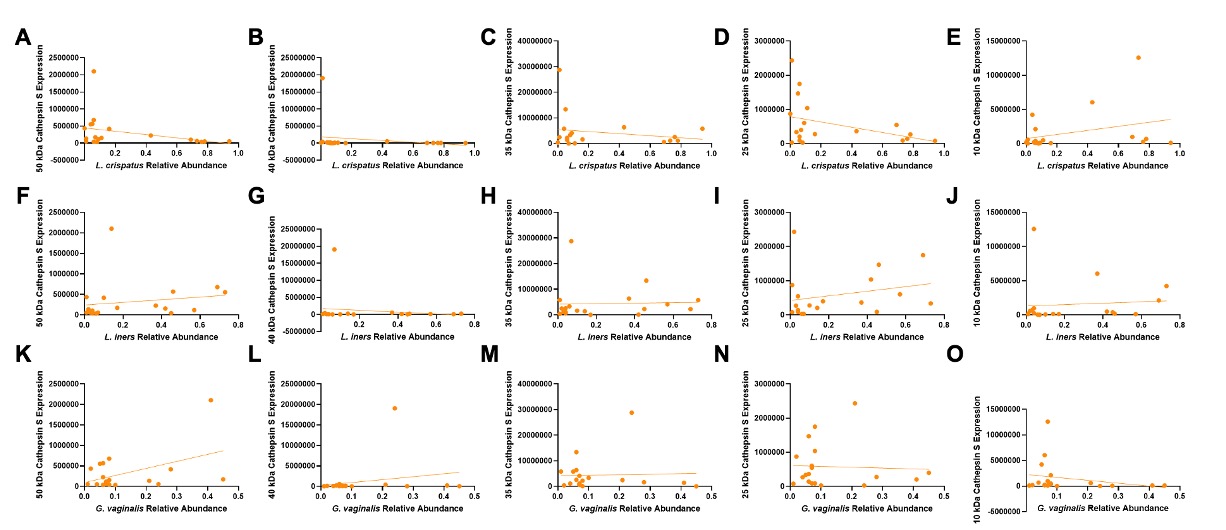

**Fig. S5** Cathepsin S expression regression analysis. By L. crispatus relative abundance, **A** prepro, **B** pro, **C** mature single, **D** mature heavy, and **E** mature light cathepsin S expression are shown; by L. iners relative abundance, **F** prepro, **G** pro, **H** mature single, **I** mature heavy, and **J** mature light cathepsin S are shown; and by G. vaginalis relative abundance, **K** prepro, **L** pro, **M** mature single, **N** mature heavy, and **O** mature light cathepsin S are shown.

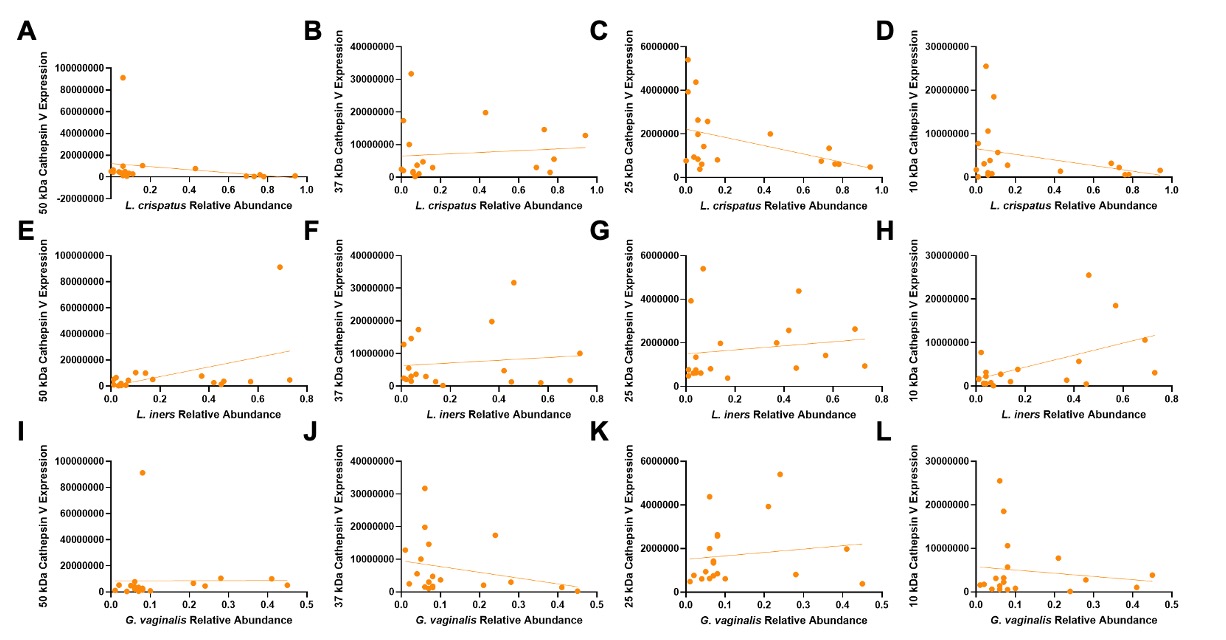

**Fig. S6** Cathepsin V expression regression analysis. By L. crispatus relative abundance, **A** pro, **B** mature single, **C** mature heavy, and **D** mature light cathepsin V expression are shown; by L. iners relative abundance, **E** pro, **F** mature single, **G** mature heavy, and **H** mature light cathepsin V are shown; and by G. vaginalis relative abundance, **I** pro, **J** mature single, **K** mature heavy, and **L** mature light cathepsin S are shown.
